# Influential factors of saliva microbiota composition

**DOI:** 10.1101/2021.12.22.473816

**Authors:** Philippa M. Wells, Daniel D. Sprockett, Ruth C E Bowyer, Yuko Kurushima, David A. Relman, Frances M.K. Williams, Claire J. Steves

## Abstract

**Background:** The oral microbiota is emerging as an influential factor of host physiology and disease state. Factors influencing oral microbiota composition have not been well characterised. In particular, there is a lack of population-based studies. We undertook a large hypothesis-free study of the saliva microbiota, considering potential influential factors of host health (frailty; diet; periodontal disease), demographics (age; sex; BMI) and sample processing (storage time), in a sample (n=679) of the TwinsUK cohort of adult twins.

**Results:** Alpha and beta diversity of the saliva microbiota was associated most strongly with frailty (alpha diversity: Q = 0.003, Observed; Q=0.002, Shannon; Q=0.003, Simpson; Beta diversity: Q = 0.002, Bray Curtis dissimilarity) and age (alpha diversity: Q=0.006, Shannon; Q=0.003, Simpson; beta diversity: Q=0.002, Bray Curtis dissimilarity; Q= 0.032, Weighted UniFrac) in multivariate models including age, frailty, sex, BMI, frailty and diet, and adjustment for multiple testing. Those with a more advanced age were more likely to be dissimilar in the saliva microbiota composition than younger participants (P = 5.125e-06, ANOVA). In subsample analyses, including consideration of periodontal disease (total n=138, periodontal disease n=66), the association with frailty remained for alpha diversity (Q=0.002, Observed ASVs; Q= 0.04 Shannon Index), but not beta diversity, whilst age was not demonstrated to associate with alpha or beta diversity in this subsample, potentially due to insufficient statistical power. Length of time that samples were stored prior to sequencing was associated with beta diversity (Q = 0.002, Bray Curtis dissimilarity). Six bacterial taxa were associated with age after adjustment for frailty and diet.

**Conclusions:** Frailty and age emerged as the most influential factors of saliva microbiota composition. Whilst frailty and age are correlates, the associations were independent of each other, suggesting that both biological and chronological ageing are key drivers of saliva microbiota composition.

## Introduction

The oral microbiota are emerging as an important factor in human disease, and their influence on host health are relatively under explored compared to those communities residing at other body sites, such as the gut. Improved understanding of these communities is imperative to appropriately design studies and determine causation in disease, with a view to develop interventions that effectively treat or even prevent disease.

The oral microbiota has previously been associated with numerous oral and systemic disease, including dental caries, periodontal disease, diseases of the oral mucosa, oral cancer and peri-implantitis [1,2], Systemic disease associated with the oral microbiota include obesity [3], rheumatoid arthritis [4], HIV infection [5], liver cirrhosis [6], inflammatory bowel disease [7], polycystic ovary syndrome [8], type 2 diabetes [9], atherosclerosis and cardiovascular disease [10] and, more recently, Alzheimer’s disease [11]. The oral microbiota are known to play an important physiological role in nitric oxide homeostasis which influences blood pressure [12].

There is an ongoing research effort to understand the composition of the oral microbiota, however previous studies have focused predominantly on disease cohorts. Studies of the general population are currently lacking. In particular, there is limited characterisation of the oral microbiota in relation to potential influencing factors such as age, sex, BMI, diet and general health (frailty). The measured composition can also be influenced by study protocol, including sample collection, storage, processing, and sequencing method. Here, we characterise the salivary microbiota present in a deeply phenotyped cohort of generally healthy adults, while accounting for demographic and protocol-related covariates.

## Methods

### Study design

The study was undertaken as an exploratory investigation of potential influential factors of saliva microbiota composition. In addition to saliva microbiota, data were collected for multiple potential contributory factors, and the aim of the study was to evaluate which of these were important with regards to saliva microbiota composition whilst controlling for the other factors.

### Participants

Participants of this study are members of the TwinsUK cohort, the largest UK registry of adult twins [13] for whom microbiome, health and dietary data had been collected as part of the ongoing data collection for the study of age-related disease. Participants are invited to visit the unit for phenotyping on a rolling basis approximately every four years. Participants enrolled in this study visited the between June 2014 and May 2017. The majority of participants were Caucasian females. The age of participants ranged from 38 to 80 (median age 66). General health status of the participants was captured using a frailty index generated from self-report diagnoses of disorders and health, following Rockwood and colleagues’ 2008 method [14]. Briefly, this is a measure of health deficit computed by dividing the number of age-associated health deficits by the total of 36 domains. Participant demographics are summarized in **Table 1**.

**Table 1.** Participant Characteristics.

|  | Age | Sex | BMI | HEI | Frailty | Zygosity | Ethnicity | No. days<br>samples stored | Fasted<br>Status |
| --- | --- | --- | --- | --- | --- | --- | --- | --- | --- |
|  | Median<br>(IQR) | N<br>(%F) | Median<br>(IQR) | Median<br>(IQR) | Median<br>(IQR) |  |  | median (IQR) | (% fasted) |
| TwinsUK<br>participants<br>(n=679) | 66<br>(15) | 625<br>(92) | 25<br>(6) | 55<br>(11) | 0.18*<br>(0.15) | MZ: 376<br>DZ: 303 | Caucasian:<br>656<br>Black: 4<br>Asian: 4<br>Mixed: 8* | 53<br>(87) | 93 |
| Dietary<br>Subsample<br>(n=582) | 67<br>(14) | 541<br>(93) | 25<br>(6) | 58<br>(12) | 0.18<br>(0.15) | MZ: 326<br>DZ: 256 | Caucasian:<br>563<br>Black: 2<br>Asian: 4<br>Mixed: 8* | 51<br>(89) | 93 |
| Periodontal<br>Disease<br>Subsample<br>(n=134) | 63<br>(15) | 134<br>(100) | 25<br>(6) | 58<br>(14) | 0.18<br>(0.16) | MZ: 77<br>DZ: 57 | Caucasian:<br>132* | 114<br>(12) | 87 |

\* Data unavailable for some participants

### Saliva sample collection

Saliva samples were collected from participants during routine volunteer visits to the NIHR BRC Clinical Research Facility associated with the Department of Twin Research at King’s College London. Participants were requested to arrive for their volunteer visit to the Clinical Research Facility having fasted for at least six hours, prior to collection of saliva samples. This was specified as abstinence from food, beverages other than water, smoking, and chewing gum for 6 hours. Participants were instructed to spit into a 30 ml sterile Falcon tube for ten minutes, and to try and produce as much saliva as possible in this time. Completed samples were immediately placed at 4 degrees Celsius before being transported in an insulated cooling bag to the laboratory within the same building. On arrival in the laboratory, samples were aliquoted into Eppendorf tubes and stored at −80°C. Frozen saliva samples were shipped on dry ice at −40°C to Stanford University for DNA extraction and 16S rRNA gene sequencing.

### Saliva sample processing and 16S rRNA gene sequencing

DNA extraction was performed using the DNeasy PowerSoil HTP 96 DNA extraction kit (Qiagen, Hilden, Germany) according to the manufacturer’s instructions. This includes a mechanical cell lysis step, of bead beating for 20 minutes. Saliva samples were randomly distributed, to avoid a twin -pair samples being placed adjacent to one another and both DNA extraction and PCR blanks were included. Prior to sequencing, samples were pooled in equimolar ratios. The V4 region of the 16S rRNA gene was amplified using PCR, in triplicate, using primers 515F and 806R which include error-correcting barcodes and illumine adaptors. Sequencing was undertaken on an Illumina HiSeq 25000 platform, generating a total of 167.8 million reads.

### Microbiota profiling

Reads were denoised using DADA2 to generate ASVs [15]. Chimeras were removed using the consensus method. Taxonomy of ASVs was assigned using the SILVA database, version 1.3.2 [16]. ASVs assigned as “mitochondria” or “chloroplast”, or which were unassigned at kingdom level were removed from the dataset. DNA extraction and PCR blanks were utilised to detect contaminant ASVs, which originate from laboratory equipment, laboratory reagents or personnel. DNA extraction and PCR blanks were used as input for the Decontam R package version 1.2.1 [17], which identified 46 contaminant ASVs. The ASVs identified as contaminants were removed from the dataset. A phylogenetic tree was generated for the ASVs by matching the ASVs into the SILVA nr v132 phylogenetic tree backbone using the fragment-insertion function (version 2018.6.17) in QIIME2.

Taxonomic and phylogenetic composition of the saliva microbiota was visualised using Phyloseq [18] and Metacoder [19] R packages, after conversion of taxon counts to per sample relative abundance.

Alpha diversity of samples was calculated from the un-trimmed and un-normalised ASV table as described in McMurdie *et al*. [20] and captured using three measures - Observed ASVs, Shannon Index and Simpson Index.

Bray-Curtis dissimilarity and Weighted Uni Frac beta diversity distances were generated after applying the negative binomial variance stabilising transformation using DEseq2 [21], Phyloseq [18] and Vegan [22] R packages.

### Statistical analysis

Analyses were undertaken using R version 4.0.3. Association of variables with alpha diversity was assessed using linear mixed effects models applied using the ‘lme4’ R package [23], and association with beta diversity was calculated using marginal permutational analysis of variance (PERMANOVA) applied via the ‘Vegan’ R package [22] with 999 permutations. Factors considered were age, sex, frailty, BMI, fasting status and length of time samples were stored in freezer. All factors were considered whilst adjusting for all other factors listed, plus sequencing depth. Dietary and periodontal disease data were not available within the full sample due to these being introduced during certain time windows of collection (data is considered missing at random). Therefore, two subsample analyses were undertaken, including participants with these data, using the same approach as above and additionally considering diet (subsample n= 582) and periodontal disease (subsample n = 134). In the periodontal disease subsample, only 4 participants were male, they were removed from the sample to avoid bias.

Differential abundance of ASVs present in more than 5% of samples was modelled against age using the ‘DESeq2’ R package, adjusting for frailty, diet, sex, BMI, whether participants had fasted prior to sample collection, storage time of samples and sequencing depth. Adjustment for multiple testing was applied to all models, using false discovery rate (FDR).

## Results

### Taxon characterisation of the saliva microbiota

Within 704 participants the saliva microbiota comprised 3,339 ASVs, comprising 15 phyla. The most dominant phyla in ascending order were Proteobacteria, Firmicutes and Bacteroidetes **(Figure 1)**.

**Figure 1.**
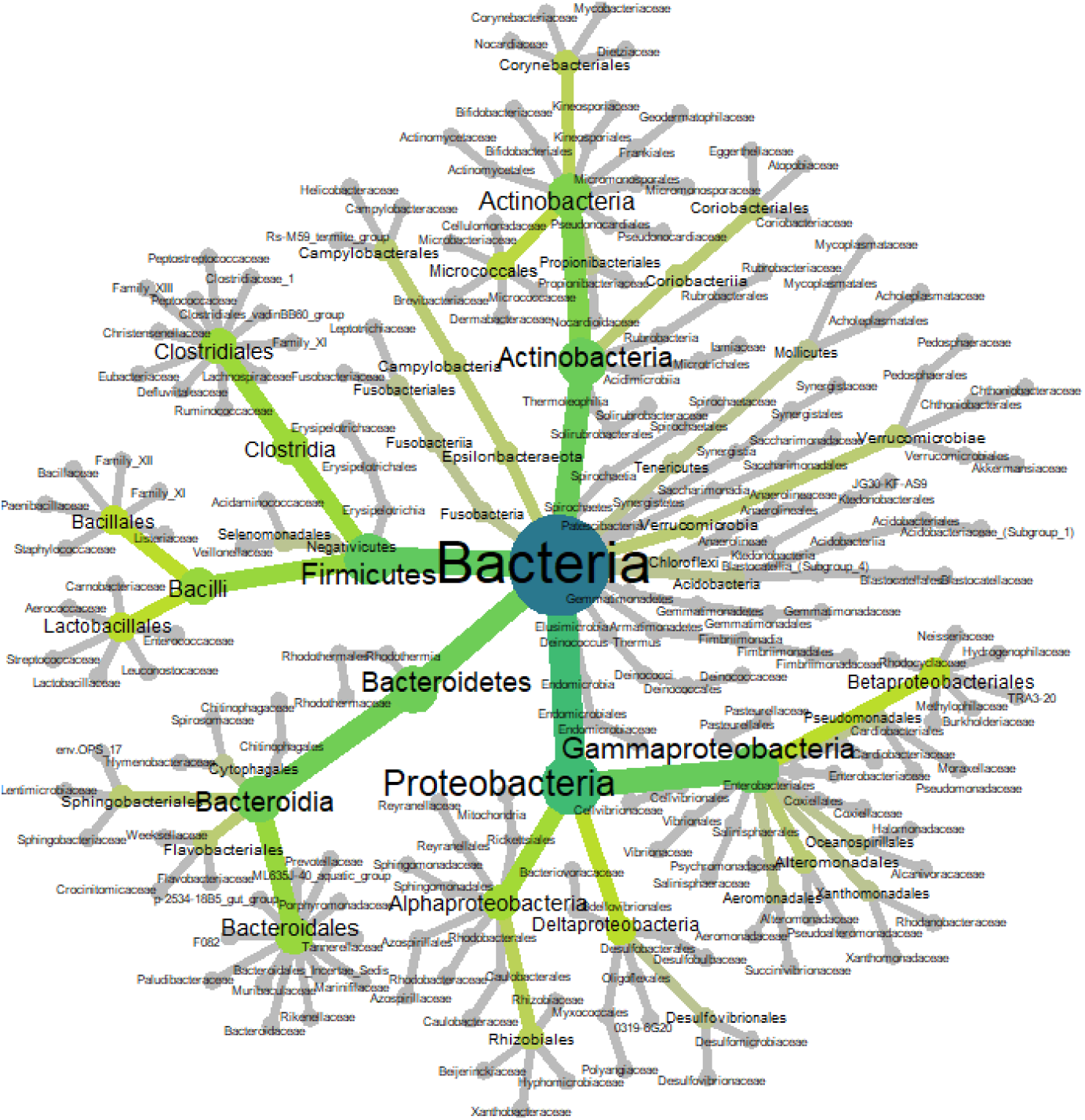
Heat tree of phylogenetic relationship and relative abundance of ASVs within the saliva microbiota of TwinsUK participants (n=679). Colour and size of nodes relate to the taxonomic level and abundance of taxa, respectively. Higher to lower taxonomic levels range from blue to dark green to light green, to grey.

The top 20 most abundant taxa across all samples comprised 63 ASVs assigned to seven phyla: Firmicutes, Bacteroidetes, Actinobacteria, Proteobacteria, Fusobacteria, Epsilonbacteraeota, and Spirochaetes. At genus assignment level, the most prevalent taxa in the cohort (defined as presence in more than 80% of samples) were *Streptococcus, Haemophilus, Veillonella, Prevotella_7, Prevotella_6, Ruminococcaceae-UCG-14, Porphyromonas, Actinomyces, Alloprevotella, Rothia, Fusobacterium, Oribacterium, Lachnoanaerobaculum, Campylobacter, Kingella, Lautropia, Cardiobacterium, Peptostreptococcus, Catonella, Mogibacterium, F0058, Bergeyella, Capnocytophagia, Granulicatella*, and *Treponema_2*.

### Diversity of the saliva microbiota

Alpha diversity was captured using three measures: Observed ASVs, Shannon diversity and Simpson Diversity. Beta diversity was captured using Bray Curtis dissimilarity and Weighted UniFrac distance. Models to investigate association with alpha and beta diversity were repeated in two sub-samples, the dietary subsample and the periodontal disease subsample with additional investigation of diet and periodontal disease, respectively.

Mixed-effects multivariate regression models were used to investigate association of the saliva microbiota with alpha diversity. An association between alpha diversity of the saliva microbiota was demonstrated for the following factors: age, frailty, sample storage time and periodontal disease **(Table 2)**.

**Table 2.** Association with saliva microbiota alpha diversity of potential influential factors. Multivariate models are presented for each sample: full sample, dietary sub-sample, and periodontal disease subsample. All models are adjusted for multiple testing and resultant Q values are given. Full sample (n=679): in a multivariate model adjusting for all other variables listed (not including diet or periodontal disease), factors associated with saliva microbiota alpha diversity were age (Q = 0.004, Shannon Index; Q=0.003, Simpson Index) and frailty (Q=0.004, Observed; Q=0.002, Shannon Index; Q=0.003, Simpson Index). In the dietary subsample (n=582), the associations demonstrated in the full sample were robust to adjustment for diet; factors associated with alpha diversity were age (Q = 0.006, Shannon Index; Q=0.003, Simpson Index) and frailty (Q=0.003, Observed; Q = 0.002, Shannon Index; Q=0.003, Simpson Index). In the periodontal disease subsample (n=134), alpha diversity was associated with frailty (Q=0.002, Observed; Q=0.04, Shannon).

|  | Full Sample |  |  |  |  |  | Dietary Subsample |  |  |  |  |  | Periodontal Disease Subsample |  |  |  |  |  |
| --- | --- | --- | --- | --- | --- | --- | --- | --- | --- | --- | --- | --- | --- | --- | --- | --- | --- | --- |
|  | Observed |  | Shannon |  | Simpson |  | Observed |  | Shannon |  | Simpson |  | Observed |  | Shannon |  | Simpson |  |
|  | *Est.* | *Q* | *Est.* | *Q* | *Est.* | *Q* | *Est.* | *Q* | *Est.* | *Q* | *Est.* | *Q* | *Est.* | *Q* | *Est.* | *Q* | *Est.* | *Q* |
| **Age** | -0.02 | 0.774 | 0.152 | 0.004** | 0.15 | 0.003** | -0.03 | 0.741 | 0.12 | 0.006** | 0.15 | 0.003** | 0.02 | 0.262 | 0.22 | 0.128 | 0.12 | 0.28 |
| **Sex (M)** | -0.03 | 0.841 | -0.293 | 0.15 | -0.26 | 0.164 | -0.01 | 0.93 | -0.28 | 0.19 | -0.26 | 0.191 | - | - | - | - | - | - |
| **BMI** | -0.02 | 0.774 | 0.06 | 0.215 | 0.08 | 0.083 | 0.02 | 0.763 | 0.06 | 0.279 | 0.08 | 0.097 | 0.18 | 0.055 | 0.14 | 0.237 | 0.12 | 0.507 |
| **Frailty** | -0.15 | 0.004** | -0.166 | 0.002** | -0.16 | 0.003** | -0.16 | 0.003** | -0.16 | 0.002** | -0.16 | 0.003** | -0.35 | 0.002** | -0.29 | 0.04** | -0.18 | 0.283 |
| **Storage Time** | 0.09 | 0.086 | 0.03 | 0.54 | 0.04 | 0.44 | 0.002 | 0.08 | 0.001 | 0.496 | 0.0001 | 0.525 | 0.01 | 0.33 | 0.01 | 0.256 | 0.005 | 0.51 |
| **Relatedness** | NA | 0.61 | NA | 0.281 | NA | 0.326 | NA | 1 | NA | 0.971 | NA | 1 | NA | 1 | NA | 1 | NA | 1 |
| **Fasting Status** | NA | 0.61 | NA | 0.281 | NA | 0.326 | NA | 1 | NA | 0.99 | NA | 1 | NA | 1 | NA | 1 | NA | 1 |
| **Diet** | - | - | - | - | - | - | -0.006 | 0.168 | -0.036 | 0.456 | 0.003 | 0.946 | -0.08 | 0.367 | 0.005 | 0.98 | 0.05 | 0.51 |
| **PD** | - | - | - | - | - | - | - | - | - | - | - | - | 0.37 | 0.055 | 0.004 | 0.981 | 0.002 | 0.991 |

Independent associations of beta diversity with age and frailty were demonstrated using a multivariate model, and this finding was robust across all models except within the periodontal disease subsample (n=134, **Table 4),** which may reflect a power issue. Frailty and the length of time that samples were stored prior to sequencing was associated with Bray Curtis dissimilarity. The association between beta diversity and age, sex, BMI, diet, frailty, participant fasting status and sample storage time is presented below **(Figure 3)**.

**Figure 3.**
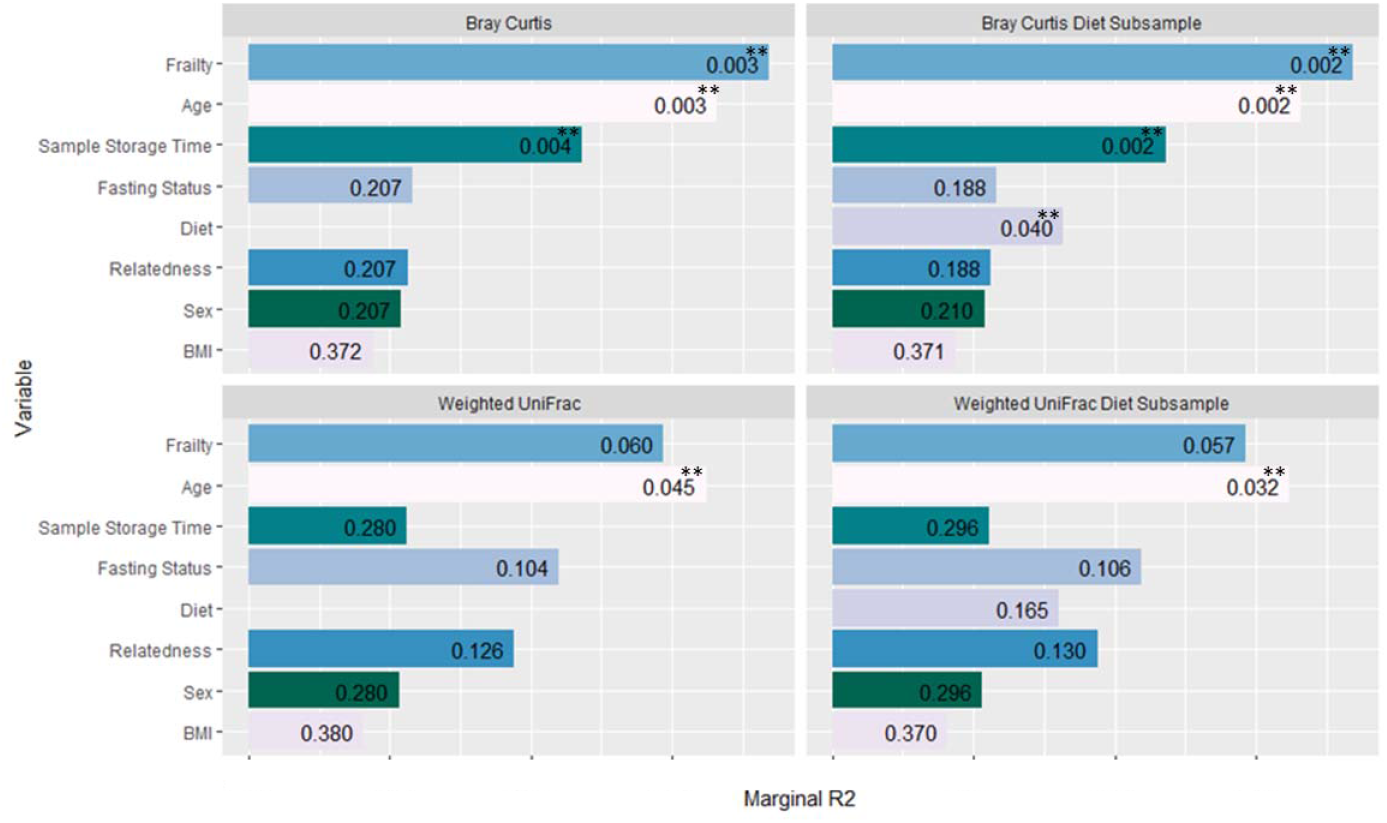
Association of factors with beta diversity of the saliva microbiota. Models using both the full sample (n=679) and diet subsample (n=582) are shown. Adonis R2 (marginal) is plotted for each variable. The diet subsample includes only participants for whom dietary data were available. Bars are annotated with FDR adjusted p values (Q values). Age was significantly associated with beta diversity after adjusting for all other factors listed plus sequencing depth when measured using both Bray Curtis and Weighted UniFrac. Frailty was associated with Bray Curtis beta diversity (Q = 0.003). Bray Curtis dissimilarity was also associated with sample storage time (length of time that samples were stored in the freezer (Q = 0.004) and diet (Q=0.04; subsample analysis).

Correlation between variables is shown in **Table 3**. A significant association was demonstrated between frailty and fasting status, whilst association between all other variables were nonsignificant.

**Table 3.**
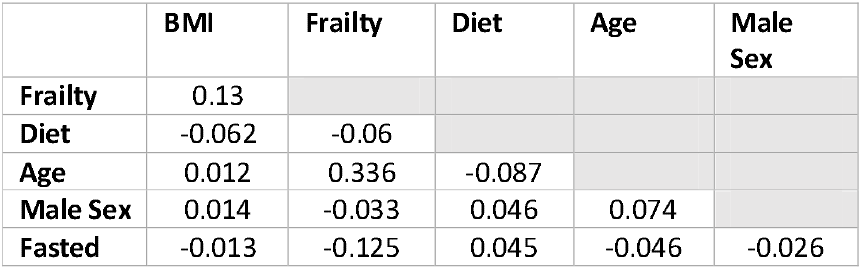
Rho coefficients of correlation between participant characteristics. Significantly correlated factors were frailty and age (p= 2.2e-16), frailty and fasting status (p=0.001), frailty and BMI (p=0.0007), BMI and having fasted prior to sample collection (p=0.001), and diet and age (p=0.035).

|  | BMI | Frailty | Diet | Age | Male Sex |
| --- | --- | --- | --- | --- | --- |
| Frailty | 0.13 |  |  |  |  |
| Diet | -0.062 | -0.06 |  |  |  |
| Age | 0.012 | 0.336 | -0.087 |  |  |
| Male Sex | 0.014 | -0.033 | 0.046 | 0.074 |  |
| Fasted | -0.013 | -0.125 | 0.045 | -0.046 | -0.026 |

**Table 4.**
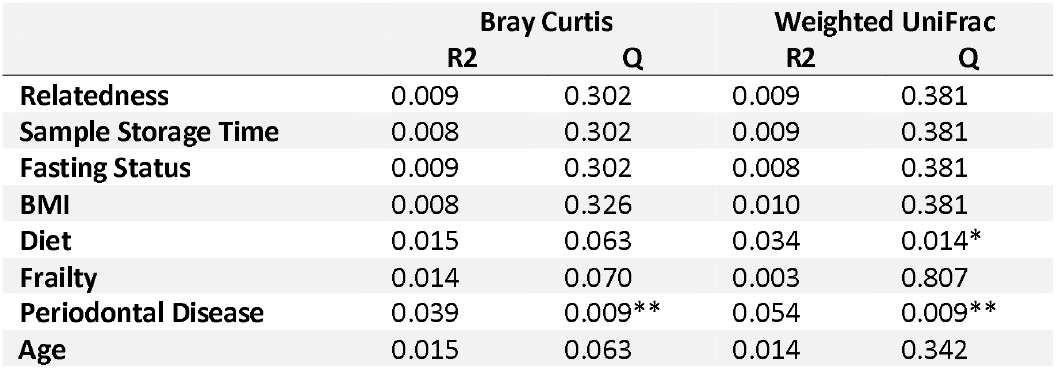
Oral health subsample PERMANOVA including participants with data for periodontal disease (n=134). In this subsample, an association of saliva microbiota beta diversity with periodontal disease (Q=0.009, Bray Curtis; Q=0.009, Weighted UniFrac) and Diet (Q=0.014, Weighted UniFrac) was demonstrated.

In the sub-sample analysis of participants with periodontal disease data, implementing the same multivariate model with the addition of periodontal disease, age was not associated with Bray Curtis dissimilarity or weighted UniFrac. This was also true for frailty and sample storage time. The loss of association of age, frailty and sample storage time with beta diversity in the periodontal disease subsample analysis may potentially be due to insufficient statistical power; the periodontal disease sub-sample was substantially smaller (n=138) than the full sample (n=679). Periodontal disease was strongly associated with both Bray Curtis dissimilarity (Q = 0.009) and Weighted Uni Frac distance (Q = 0.009).

Across 5 age groups, variation in distribution of beta diversity increased sequentially with age, demonstrating that younger participants were more similar in their saliva microbiota diversity than older participants **(Figure 3)**.

**Figure 3.**
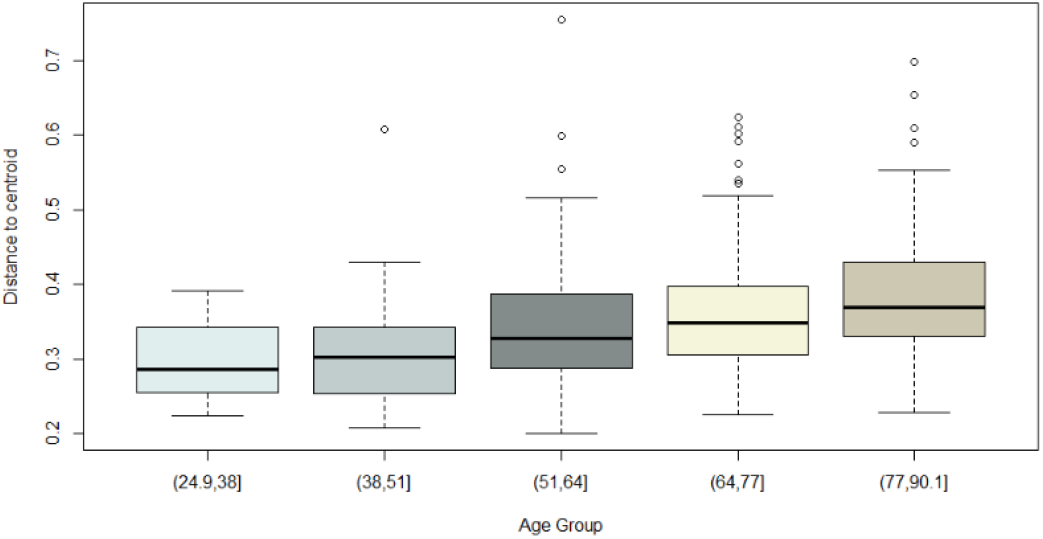
Bray Curtis beta dispersion of age within full cohort sample. A difference in the variance across age groups was demonstrated (P = 5.125e-06, ANOVA).

### Saliva microbiota taxa associated with age

The association of saliva microbiota composition age was explored taxonomically. After adjustment for all factors available for the dietary data subsample (n=582, **methods),** and multiple testing, six ASVs were associated with age **(Figure 4)**.

**Figure 4.**
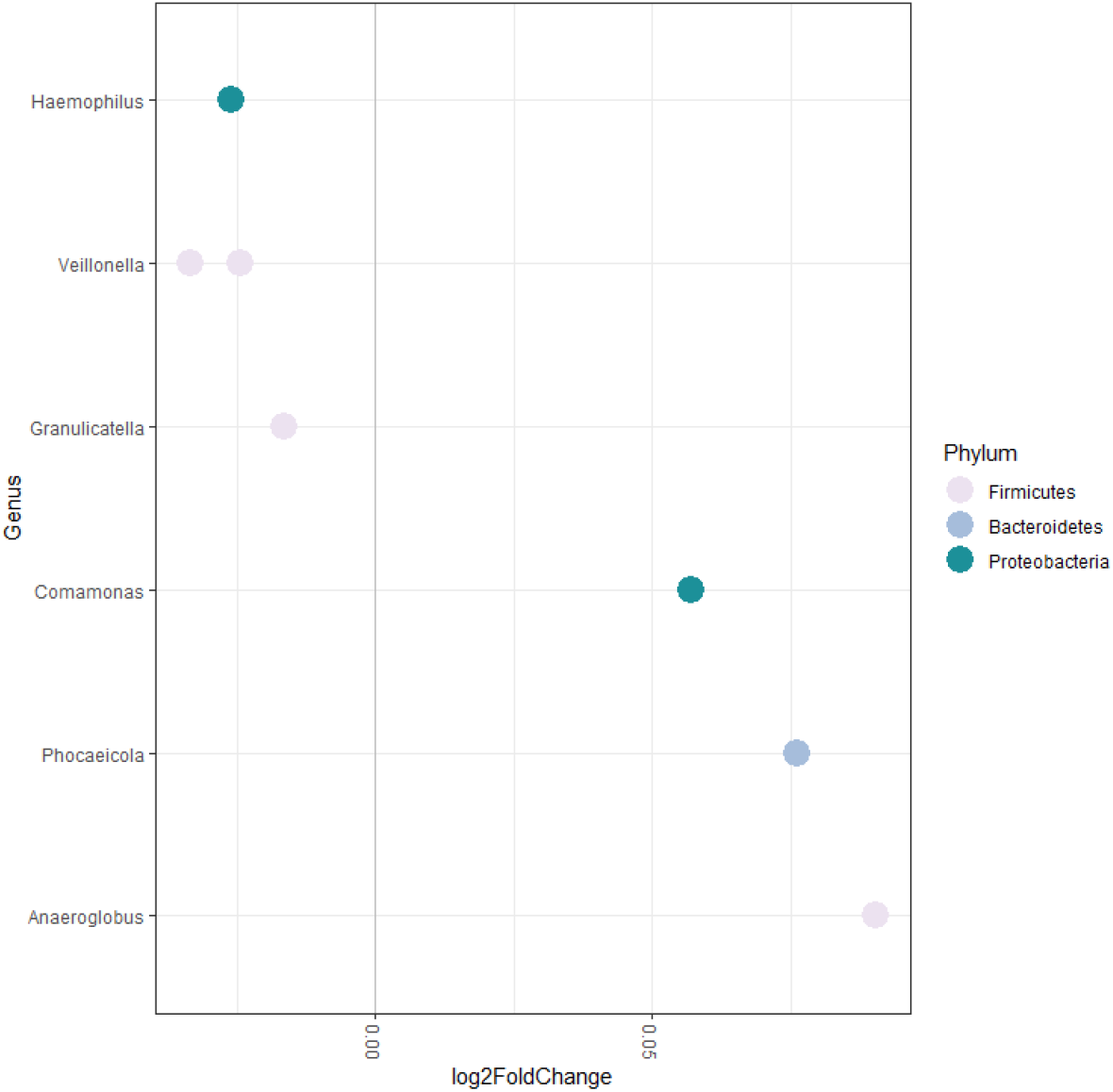
Saliva microbiota associated with age. Age was associated with six taxa in a multivariate model after adjustment for sex, frailty and diet of participants. Taxa positively associated with age were *Comamonas* (Q= 0.029), *Phocaeicola abscessus* (Q = 0.037) and *Anaeroglobulus germinatus* (Q = 0.004). Taxa inversely associated with age were *Veillonella* (Q = 0.0002), *Haemophilus* (Q = 0.0004), *Veillonella atypica/dispar* (Q = 0.037) and *Granulicatella adiacens/para-adiecens* (Q = 0.05).

## Discussion

The composition of the saliva microbiota requires characterisation in order to understand links with health and disease. In this explorative study we performed a multifactorial investigation of the saliva microbiota in a large, unselected, population sample from TwinsUK. Age and frailty (health deficit) emerged independently from each other as the strongest influential factors of alpha and beta diversity, after adjustment for all other factors. Taxonomically the association with age was driven by six bacterial species.

These findings suggest that both chronological and biological ageing may be important with regards to saliva microbiota composition. Potentially, there may be an influence of the ageing immune system and immunosenescence, which is also thought to underly increased incidence and severity of infection with increasing biological age (frailty) [24].

There are limited prior studies of the saliva microbiota composition, however these have demonstrated an association of diversity with BMI [25] and frailty [26]. In a study by Wu *et al*. which characterised the salivary microbiome of 62 younger adults in relation to obesity, an association of BMI was demonstrated with both alpha diversity (Chao1, P < 0.01; Shannon diversity, P < 0.05) and beta diversity differences (unweighted UniFrac, P = 0.001) [25]. The study design accounted for age, sex and oral hygiene. In contrast, in TwinsUK, no association for BMI was demonstrated, although our sample comprised older adults with lower average BMI (25) and modest intra sample variation (SD 6). In addition, Wu *et al*. targeted the V3 variable region, which could potentially account for the difference in results if there is higher sensitivity for taxa which associate with obesity.

A study of the saliva microbiota by Ogawa *et al*. included consideration of frailty. Of participants, 16 lived in a nursing home and 15 lived within the general community [26]. Those who were nursing home dwellers had a mean age of 87, whereas those who lived within the community had a mean age of 84. Participants were admitted to the nursing home on recommendation by a medical doctor due to frailty. The authors therefore classified the nursing home group as frail and the independent community dwellers as non-frail. Ogawa *et al*. demonstrated a significant inverse association of alpha diversity of the saliva microbiota with nursing home dwelling, which was interpreted to be due to frailty. They accounted for age, BMI and dental health, but included no adjustment for diet. A difference in diet between the non-frail and frail groups could be an important unmeasured confounder, given the dwelling of participants. Our study was conducted in community dwelling older adults, and measured frailty using the frailty index. We did adjust for diet as well as other covariates and found a significant inverse association of frailty with alpha diversity, this corroborating their results. Additionally, in the present study we found evidence that frailty was associated with different microbiota composition (Bray Curtis dissimilarity and Weighted UniFrac distance).

A recent population study by Burcham *et al*., considered factors which influence the oral microbiota in adults versus children (Burcham et al. 2020). Oral samples were obtained using buccal swabs of the teeth, tongue, cheeks and gums. Therefore, multiple oral sites were included, in addition to saliva. The source of saliva microbiota is predominantly the biofilm on the dorsum of the tongue, however all other oral microbial niches contribute taxa (Davenport 2017). The buccal swabs used by Burcham are therefore relevant to the saliva microbiota but may not be directly comparable. Burcham *et al*. included 172 adults aged between 20 and 57, median age 34. They did not consider age within the adult or child groups, but on comparison of both groups they demonstrated significantly lower Shannon diversity in adults versus children. Within both groups they considered weight status, sex, prescription of antibiotics in the last 6 months and oral hygiene habits (visits to the dentist for professional descaling). They showed that in adults only, composition of the microbiota (beta diversity) varied with oral hygiene habits. There was no association between weight status, sex or ingestion of antibiotics in last 6 months. However, youth oral microbiome beta diversity was associated with both sex and weight (Burcham et al. 2020). In the present study, oral hygiene habits were not accounted for, but could potentially be explain some of the signal identified for age and frailty.

In a study by Zhang *et al*, in which they considered age, sex, BMI and inflammatory and rheumatoid arthritis disease markers, there was no association of any factors with beta diversity of the saliva microbiota [27]. However, age was the only factor which was close to significance, and study was limited by small sample size (n=98).

A technical factor - length of time that samples were stored prior to sequencing - also emerged as a distinguishing feature of beta diversity, suggesting that some taxa are more affected than others by long-term freezing. This will be relevant to many studies using larger cohorts which collect data over a wide time span [28].

Moving forward, it will be important to investigate the role of host genetic factors in determining the composition of the saliva microbiota. A recent study using twin based heritability estimates of saliva microbiota from 209 twin pairs found that whilst overall non-shared environment is the most important factor [28,29], substantial heritability was demonstrated for 28 percent of ASVs in their samples [28]. They also demonstrated heritability of differential immune response to commensal oral microbes, which would provide a mechanism for heritability of the microbiota.

In TwinsUK saliva samples, the most abundant phylum within the saliva microbiota was demonstrated to be Proteobacteria **(Figure 1)**. In our study, employing sequencing of the V4 16SrRNA gene variable region, the taxonomic composition demonstrated is in contrast with prior studies, the majority show Bacteroidetes to be the dominant phylum [26,30–32]. In these prior studies, a key common difference accounting for taxonomic discrepancy is choice of primer; these prior studies undertook sequencing of the V2 variable region. Murugesan *et al*. undertook a large recent study of the saliva microbiota of 997 younger adults with mean age of 38, using targeting of the V2 variable region, demonstrated that the dominant phylum of the saliva microbiota was Bacteroidetes [32]. Proteobacteria were the third most dominant phylum. This finding was replicated in studies by Ogawa *et al*., *Gomez et al*., and Tsuda *et al*. [26,30,31].

In the study by Ogawa *et al*. of 31 elderly adults, within community dwelling older adults, the saliva microbiota were dominated at the phylum level by Bacteroidetes. Proteobacteria was the third most abundant phylum, after Firmicutes. There are important methodological differences compared to the present study, which are likely to account for the taxonomic discrepancy. The most influential methodological discrepancy is likely to be choice of primers. However, in addition they generated *de novo* operational taxonomic units (OTUs) and assigned taxonomy of using the Greengenes database. In the present study ASVs were generated and taxonomy was assigned using the SILVA database. Ogawa and colleagues required their participants to refrain from food, drink, smoking and use of toothpaste for 2 hours prior to saliva collection. In the present study, participants were asked to fast and refrained from smoking or chewing gum for at least 6 hours, whereas there was no stipulation about toothpaste. Ogawa *et al*. undertook DNA extraction, PCR and sequencing immediately, whereas in the present study samples were frozen prior to these steps. The primary factor mediating the taxonomic discrepancy between both studies is likely to be choice of variable region targeted by the PCR primers.

Tsuda *et al*. undertook a study of the saliva microbiota of 44 adults who had fasted overnight, using pyrosequencing after PCR targeting of the V1-V2 hypervariable region [30]. The dominant phylum was Firmicutes, whilst Bacteroidetes, Actinobacteria and Proteobacteria were the second, third and fourth most abundant, respectively. Another study, by Lundmark *et al*. evaluated the saliva microbiota of 114 adults, using targeting of the V3-V4 hypervariable region [33]. They demonstrated that the most abundant phyla in order of dominance were Firmicutes, Bacteroidetes, and Proteobacteria.

There are few studies of the saliva microbiota which use targeting of the V4 variable region, however in a small study of 20 adult participants, using this method, in accordance with our study, Proteobacteria was the most dominant phylum, whilst Firmicutes was the second most dominant [34]. In the study by Gomez *et al*. the dominant phylum of the subgingival plaque of children aged 5 to 11 is Firmicutes [31]. However their study was not directly comparable, being of children, and using primers which target the V2 variable region rather than V4.

These high phylogenetic level taxon discrepancies between studies which employ targeting of different variable regions of the 16S rRNA gene indicate that primer selection is an important factor having influence on downstream findings. The primers used in this study, 515F and 806R, were initially used by the Earth Microbiome Project, and have since been modified for use with the Illumina platform [35]. This modification rectified the previously held bias against *Crenarchaeota* and *Thaumarchaeota* [36]. These primers have been demonstrated to perform well for characterising the gut microbiota, however efficacy/performance for characterisation of the oral microbiota requires further investigation. A recent study showed a discrepancy in performance when applied to human gut versus skin microbiota [37]. Most studies of the saliva microbiota have employed primers which target the V2 variable region. This region demonstrated higher resolution for *Streptococcus*, the most abundant genus in the oral cavity [38].

Our study benefitted from several strengths. Firstly, this was a large, relatively unselected population of older community dwelling adults with a breadth of data which enabled us to consider a number of potentially important factors, including host general health, diet and periodontal disease. There are key strengths in the technical aspects of sample processing: DNA extraction and sequencing was performed by the same person, samples from twin pairs were separated on the DNA extraction plates and all samples were included in two lanes of the same sequencing run. The inclusion of blank reagent only samples allowed for removal of potential contaminant ASVs from the dataset, sometimes referred to as the “kitome”. Consideration of contaminants may be particularly important for lower biomass samples such as saliva. There were also limitations to the work which are important to consider. Our sample is a volunteer cohort and there may therefore be a healthy volunteer bias. For example, in relation to frailty, our sample in accordance with the wider TwinsUK cohort is comprised of relatively healthy (lower frailty), community dwelling participants, and may not reflect changes at high levels of frailty. Similarly, participants are older adults, but few participants were aged over 70 or less than 40. Despite this, however we were able to demonstrate importance of these factors. A possible limitation was that we used targeting of the V4 region in order to analyse concurrently with gut microbiota, which, as discussed above, may have lower resolution for key taxa of the oral microbiota, and particularly *Streptococcus. We* used sampling of the saliva microbiota, which derive predominantly from oral mucosal surfaces. However, the microbiota of other sites, and particularly subgingival plaque may be more pathologically relevant. In our study we did not include an investigation of oral hygiene or smoking habits.

## Conclusions

In a multivariate exploratory study of a population sample, TwinsUK, composition of the saliva microbiota was associated with age and frailty, indicating that both chronological and biological ageing are implicated. Diversity of the saliva microbiota increased with age; younger participants were more likely to have a similar saliva microbiota composition, whereas older participants demonstrated wider difference. Storage time before processing was a key methodological variable. There were six bacterial taxa within the saliva microbiota associated with age. Our findings highlight the importance of time (both in vivo and ex vivo) and general health in the composition of the salivary microbiota composition and highlight the need to take account of these factors in any study of the association of the salivary microbiota and disease.

## Declarations

### Ethics approval and consent to participate

Ethics approval was granted by the St. Thomas’ Hospital Research Ethics Committee. Following the restructure and merging of the research ethics committee, subsequent amendments were approved by the National Research Ethics Service (NRES) Committee London–Westminster (TwinsUK reference EC04/015); approval for the use of microbiota samples was granted by the NRES Committee London–Westminster (The Flora Twin Study reference 12/L0/0227).

### Availability of data and material

The data utilised in this study is available upon reasonable request to TwinsUK.

### Competing interests

The authors declare no competing interests

### Funding

This work was funded by a Versus Arthritis Special Strategic Award (grant 21227). Department of Twin Research & Genetic Epidemiology at King’s College London receives grant support from the Wellcome Trust (212904/Z/18/Z), the MRC British Heart Foundation Ancestry and Biological Informative Markers for Stratification of Hypertension (AIMHY; MR/M016560/1), the European Union, CDRF, ZOE Global, NIH, the National Institute for Health Research (NIHR)-funded BioResource, and Clinical Research Facility and Biomedical Research Centre based at Guy’s and St Thomas’ National Health Service (NHS) Foundation Trust, in partnership with King’s College London. The funding providers for this work did not provide input regarding study design

### Authors Contributions

CJS conceived the study, DS sequenced the saliva samples and generated the ASVs, DR supervised the work of DS. PW analysed the data and drafted the manuscript. RB and YK contributed to the analysis methods. CJS and FMKW supervised the work of PW.

## Acknowledgements

The authors would like to thank the participants of the TwinsUK cohort.

## Notes

### Competing Interest Statement

The authors have declared no competing interest.

